# A Micro-engineered Human Colon Intestine-Chip Platform to Study Leaky Barrier

**DOI:** 10.1101/2020.08.28.271759

**Authors:** Athanasia Apostolou, Rohit A. Panchakshari, Antara Banerjee, Dimitris V. Manatakis, Maria D. Paraskevopoulou, Raymond Luc, Galeb AbuAli, Alexandra Dimitriou, Carolina Lucchesi, Gauri Kulkarni, Tengku Ibrahim Maulana, Bertram Bleck, Elias S. Manolakos, Geraldine A. Hamilton, Cosmas Giallourakis, Katia Karalis

**Affiliations:** Emulate Inc., 27 Drydock Avenue, Boston, MA, 02210, USA; Graduate Program, Department of Medicine, School of Health Sciences, National and Kapodistrian University of Athens, Athens, 11527, GR; Takeda Pharmaceuticals, Cambridge, MA, 02139, USA; Takeda California Inc., San Diego, CA, 92121, USA; Department of Informatics and Telecommunications, National and Kapodistrian University of Athens, Athens, 16122, GR; Northeastern University, Boston, MA, 02115, USA; Graduate Program, Faculty of Energy-, Process- and Bioengineering, Department of Bioengineering, University of Stuttgart, Stuttgart, 70569, DE

## Abstract

The intestinal epithelial barrier supports the symbiotic relationship between the microbiota colonizing the intestinal epithelium and the host immune system to maintain homeostasis. Leaky barrier is increasingly recognized as part of the pathogenesis of a number of chronic conditions in addition to inflammatory and infectious diseases. As our understanding on the regulation of the barrier remains limited, effective therapeutic targeting for the compromised barrier is still an unmet need. Here we combined advancements on the organoids and Organ-on-Chip technologies to establish a micro-engineered Colon Intestine-Chip for studying development and regulation of the human intestinal barrier. Our data demonstrate the significance of the endothelium in co-culture with the epithelial cells within a tissue-relevant microenvironment for the establishment of a tight epithelial barrier of polarized cells. Pathway analysis of the RNA sequencing (RNA-Seq), revealed significant upregulation of mechanisms relevant to the maturation of the intestinal epithelium in organoid-derived epithelial cells in co-culture with endothelium as compared to organoids maintained in suspension. We provide evidence that the Colon Intestine-Chip platform responds to interferon gamma (IFNγ), a prototype cytokine utilized to model inflammation-induced barrier disruption, by induction of apoptosis and reorganization of the apical junctional complexes as shown with other systems. We also describe the mechanism of action of interleukin 22 (IL-22) on mature, organoid-derived intestinal epithelial cells that is consistent with barrier disruption. Overall we propose the Colon Intestine-Chip as a promising human organoid-derived platform to decipher mechanisms driving the development of leaky gut in patients and enable their translation for this unmet medical need.

## INTRODUCTION

The gastrointestinal barrier is the most extensive epithelial barrier of the body that is colonized by commensal bacteria, segregated from the host’s immune system to maintain homeostasis^1^. The characteristics of the intestinal barrier are mainly attributed to the epithelial cell interconnections via specialized apically located tight junctions (TJs). Supporting elements include locally secreted factors, the covering mucin layer, and humoral factors derived from the microbiome. Disruption of the barrier has been first identified in gastrointestinal infections and inflammatory bowel disease (IBD)^2, 3^; while the list is continuously expanding to include additional diseases, drugs and toxic factors^4, 5^. Intestinal crypt stem cells continuously renew intestinal epithelial cells (IECs) during the lifespan, thus maintaining a physiological barrier in homeostasis and enabling the repair of injured barrier in a pathophysiological state. Dysregulation of intestinal epithelial regeneration is thought to be involved in the establishment of chronic GI diseases.

The significance of cytokines in disruption, repair, or accelerated damage of the intestinal barrier has been demonstrated both *in vivo* and in cell-based disease models. The mechanisms that the proinflammatory cytokines employ to disrupt the barrier include induction of epithelial cell apoptosis and degeneration of the tight junctions. IFNγ is the prototype cytokine used experimentally, either alone or in combination with tumor necrosis factor alpha (TNFα) or interleukin-1 beta (IL-1β), to induce barrier disruption with reproducible effects in several different intestinal models. Prolonged exposure of IECs to IFNγ results in activation of apoptotic mechanisms^6^, increased expression of dickkopf WNT signaling pathway inhibitor 1 (DKK1), and inhibition of epithelial proliferation^7^, as well as phosphorylation of myosin light-chain (MLC) via activation of the MLC kinase (MLCK), that drives internalization and thus, disruption of TJs^8, 9^.

A major advancement in our ability to understand the regulation of the barrier and its actual contribution in human diseases has been the discovery of organoid technology^10–12^. This tremendous development not only enabled the use of patient-derived samples that allow the spectrum of normal and pathological individual responses ^13^, but also advanced significantly our understanding on the intestinal epithelial biology leveraging human primary cell-based pharmacological models. Recently, a culture system enabling the expansion of organoids to monolayer multicellular polarized epithelial cultures has been developed in transwells and applied to model intestinal infection^14^. This system bypasses a main caveat of the organoids, the accessibility to the apical side of the cell, the equivalent of the luminal tissue side. A missing element of all the above systems remains the ability to incorporate conditions in culture that emulate the intestinal microenvironment. The Organ-on-Chip technology has been developed to close this gap by incorporating key features of the tissue microenvironment^15–17^. Specifically, for the gut, significant features include stretching to emulate the peristalsis-like motion of the tissue, exposure to shear stress and interactions of the epithelium and endothelium, or other cells via tissue-relevant extracellular matrix proteins. We have recently demonstrated that the Duodenum Intestine-Chip supports the synchronous culture of different cell types that build up the intestinal wall and enable functionality much closer to the in vivo states compared to other systems^18^.

In this work, we leveraged the Organ-on-Chip technology^17, 19, 20^ and the proprietary Human Emulation System, as previously described, to develop a Colon Intestine-Chip platform based on human organoid-derived epithelial cells and human primary microvascular endothelial cells. First we established an optimized, *in vivo* relevant cytoarchitecture, with tissue-relative abundance of the key cell subtypes and tight barrier functionality and demonstrated the reproducibility in several donors. Next, we employed the Colon Intestine-Chip to establish a model of IFNγ elicited barrier disruption and have confirmed its described mechanism of action. Finally, we used the Colon Intestine-Chip to examine the effect of IL-22, a cytokine released by immune cells and acting on IECs, with conflicting data about its role on epithelial integrity. We show that IL-22, upon engaging its receptors expressed in the Colon Intestine-Chip, acts as a barrier-insulting cytokine, in line with recent publications^21–23^. These findings when considered together indicate that the Colon Intestine-Chip renders a promising human platform for studying intestinal epithelial development and homeostasis.

In summary, we present the development of a new biopsy derived Colon Intestine-Chip that provides a human organoid-based platform to study cell-cell interactions and mechanisms driving barrier disruption by cytokines. The Colon Intestine-Chip is a significant development as leaky barrier remains an unmet medical need, even though it is a process implicated in the pathogenesis of the majority of gastrointestinal diseases, toxicity of frequently used drugs, and other processes such as aging.

## RESULTS

### Development of a Colon Intestine-Chip platform

It has been previously described that the juxtaposed culture of intestinal epithelial and endothelial cells in micro-engineered Organ-Chips, results in the formation of a three-dimensional mucosal layer^17, 24^. Here, we applied this technology to a Colon Intestine-Chip based on enzymatically fragmented human organoids isolated from colonic biopsies, along with colon-specific microvascular endothelial cells (Figure 1A, C). The fragmented organoids were introduced in the top channel of the chip and seeded on a membrane uniformly coated with extracellular matrix (ECM) proteins, as confirmed by fluorescent staining with N-Hydroxysuccinimide (NHS)-ester dye (Figure 1B). The fluidic culture was performed using the semi-automated Human Emulation System called Zoë (Emulate, Inc). Culture conditions included fluid flow at 60 μL/h initiated on Day 1, followed by the removal of the stemness supporting factors, Y-27632 and CHIR99021, on Day 2 of the culture (Figure 1C, D). Periodic stretch was inititated on Day 3 at 2% intensity and following 24 h pre-acclimation, was switched to 10% for remainder of the study. The key point for switching to 10% stretching is when the apparent permeability of the epithelial barrier to 3kDa Dextran is stabilized below 0.5 × 10^-6^ cm/s^25^ (Figure 2B, Supplementary Figure 1B). Twenty-four hours after the initiation of 10% stretching, we assessed the formation of tight junctions in the expanded colonic epithelial monolayer by immunostaining for ZO-1 and E-cadherin (Figure 2C, Supplementary Figure 1C). In addition, we evaluated the polarized expression of ion channels by immunostaining for the apically expressed DRA, and for the basolaterally expressed Na^+^/K^+^ATPase transporter (Figure 2D, Supplementary Figure 1D). These results demonstrate the establishment of a tight, polarized, human colonic epithelial monolayer by Day 5 of the culture of colonic organoids on the Intestine-Chip.

**Figure 1.**
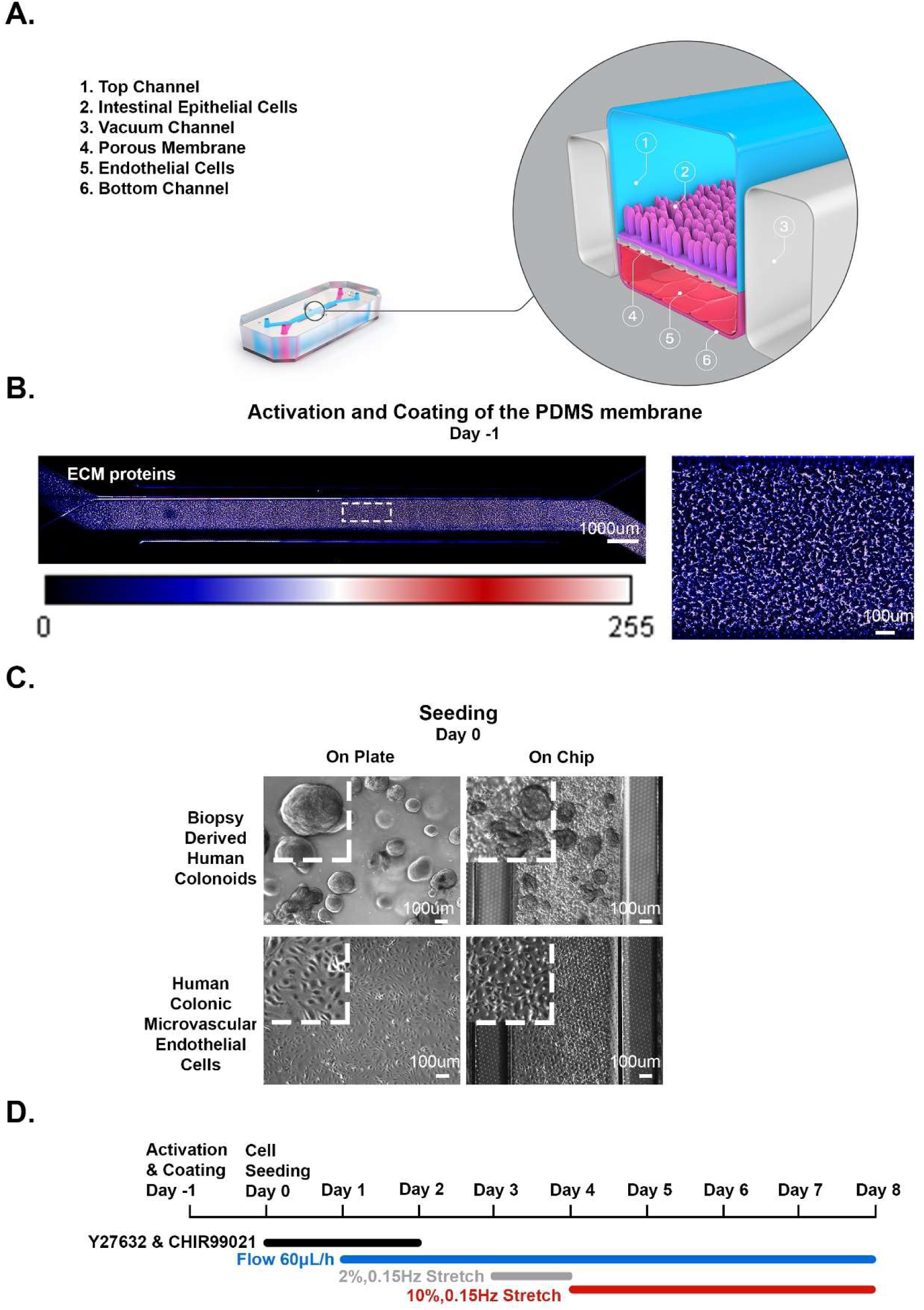
Establishment of the Colon Intestine-Chip. **[A]** Cross section of the Colon Intestine-Chip showing: the epithelial channel (1), colonic IECs (2), lateral vacuum chambers (3), porous PDMS membrane (4), cHIMECs (5) and endothelial channel (6). **[B]** NHS Ester Dye staining against the ECM proteins, Collagen IV and Matrigel, of the apical channel and the respecting LUT color bar, representing the values of an 8-bit image pixel intensity. **[C]** Representative contrast phase microscopy images of each cell type morphology, on a conventional plate culture (left) and after their introduction in the Colon Intestine-Chip (right), on Day 0 of the culture. Scale bar: 100μm **[D]** Experimental timeline of the 8 days fluidic culture.

**Figure 2.**
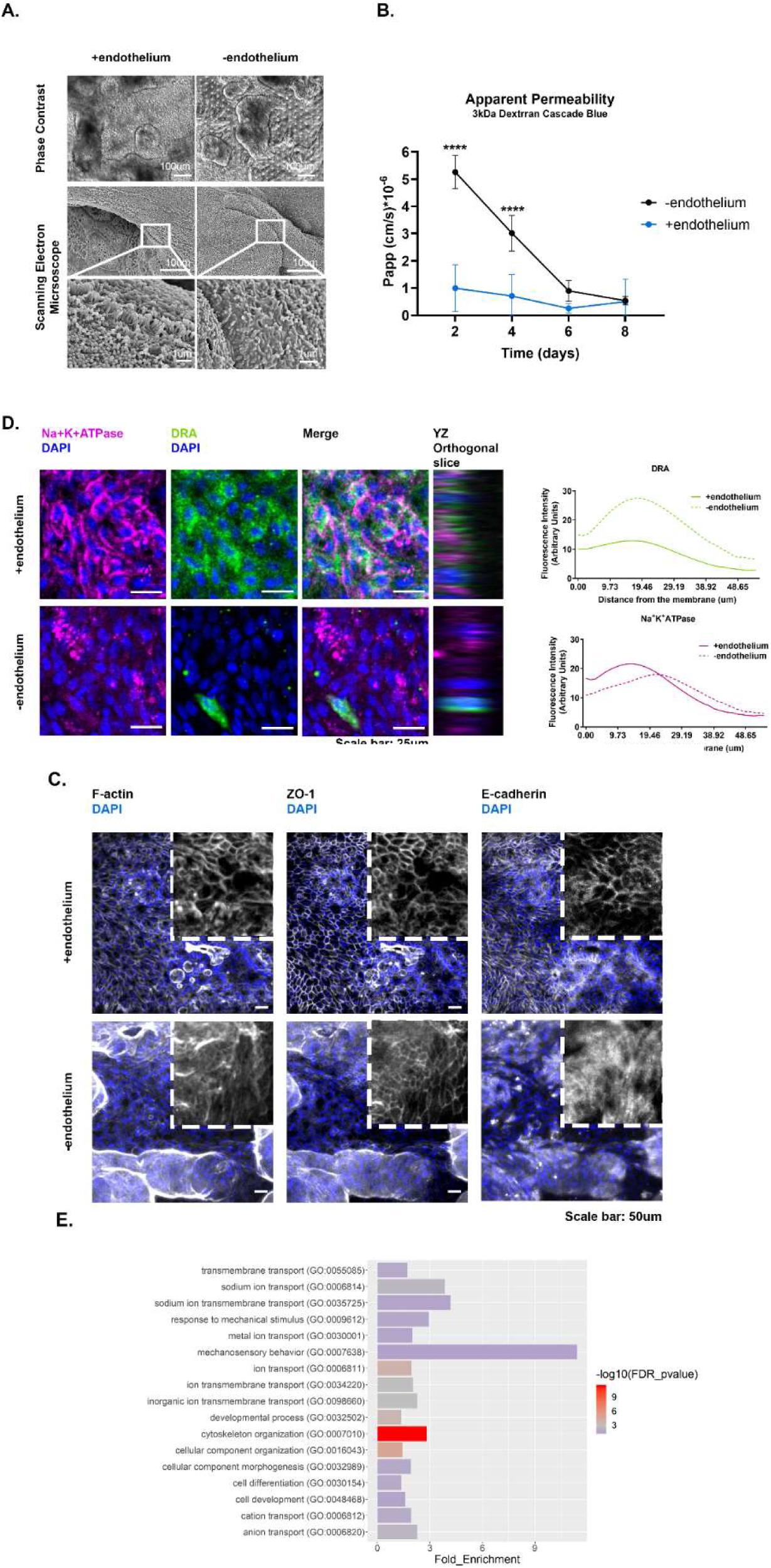
The presence of endothelium enhances the establishment of the epithelial barrier. **[A]** Representative phase contrast and Scanning Electron Microscopy (SEM) images of the colonic epithelial on day 5 of the Colon Intestine-Chip culture indicating enhanced maturation of the epithelial brush border in the presence of endothelial cells. **[B]** Apparent Permeability (P_app_) of the colonic epithelial cells to 3kDa Dextran Cascade blue, over the course of an 8 day culture, where stretch (10% strain, 0.15Hz) was applied in the presence or absence of endothelium. Data shown correspond to one representative out of three independent experiments. n= 3-11 chips/condition, mean±95% CI, Two-way ANOVA, Tukey’s post hoc test, ****: p<0.0001. **[C]** Representative confocal fluorescent images showing the establishment of strong epithelial tight junctions on day 5 of the Colon Intestine-Chip culture, in the presence of endothelium. Tight junctions were stained with anti-ZO-1 and anti-E-cadherin and cytoskeleton with phalloidin, all shown in gray. Cells nuclei are shown in blue. Scale bar: 50 μm. **[D]** Representative confocal immunofluorescence images indicating the establishment of a polarized epithelial monolayer only during the co-culture of colonocytes with endothelium. The basolateral transporter Na^+^K^+^ATPase is visualized in magenta and the apical transporter DRA in green. Cells nuclei are shown in blue. The plots indicate the mean fluorescent intensity distribution for each channel, DRA (green) or Na^+^K^+^ATPase (magenta), along the basal-apical axis of the epithelial cells. Three FOVs were acquired per chip. n= 3 chips/ condition. Scale bar: 25 μm **[E]** Gene Ontology biological processes analysis, comparing epithelial cells on day 5 of culture, where stretch (10% strain, 0.15Hz) is applied in the presence or absence of endothelium, indicates the upregulation of terms related to the morphogenesis of the colonic epithelial barrier.

We have maintained the Colon Intestine-Chip in culture for up to 12 days and the epithelial barrier integrity was monitored on a daily basis. We show here data from cultures maintained for 8 days, with reproducible and robust data across different donors.

### Role of the endothelium in the establishment of the epithelial barrier

In the Colon Intestine-Chips when organoids were co-cultured with or without endothelial cells, as described in Methods, we observed that the presence of endothelial cells enhanced epithelial barrier integrity, as indicated by the apparent permeability to 3kDa Dextran (Day 4, minus endothelium: 3.01 × 10^-6^ cm/s versus plus endothelium: 0.89 × 10^-6^ cm/s) (Figure 2B, Supplementary Figure 1B). In line with this observation, we identified a mature network of TJs as early as in the Day 5 of culture (Figure 2C, Supplementary Figure 1C). On the contrary, in the absence of endothelium the TJs were poorly defined, and ZO-1 and E-cadherin staining detected the proteins localized in the cytoplasm, rather than the membrane of the cells, as anticipated in the presence of a tight epithelial barrier (Figure 2C). These data suggest that the endothelium supports the establishment and the maturation/differentiation of the intestinal epithelium and highlights the advantages of co-culture systems. Further, immunostaining against the basolateral transporter Na^+^/K^+^ATPase and the apical transporter DRA, highlighted the compromised formation of epithelial cell polarity in the absence of endothelium (Figure 2D). On the other hand, application of stretching did not have a measurable impact in the polarization of epithelial cells (Supplementary Figure 1D). Parallel Scanning Electron Microscopy analysis revealed the formation of a mature brush border, with densely packed microvilli, only in the presence of endothelium (Figure 2A).

To comprehensively characterize the co-culture effects of colonoids and endothelial cells on the chip at a transcriptome wide level, we performed RNA-Seq analysis of the colonoids cultured in suspension and the colonoid-derived epithelial cells on-Chip, in the presence or absence of endothelium and/or stretch. Principal Component Analysis, employed on the highly variable genes across libraries, showed distinct gene expression profiles between the different on-Chip culture conditions and the colonoids in suspension. Our data highlights the effect of endothelium on the divergence of the transcriptome of colonic epithelial cells cultured on-Chip as opposed to colonoids in suspension (Supplementary Figure 1G, H). We performed functional analysis on the 573 differentially expressed genes, identified out of the 58,342 genes annotated in the genome between IECs cultured with or without endothelium in the presence of stretch, on Day 5 of culture (Supplementary Figure 1E, F). Significantly enhanced terms included terms related to cell development and morphogenesis, cytoskeletal remodeling, mechanosensory behavior and membrane ion transportation (Figure 2E). These findings are consistent with the aforementioned experimental readouts, as for example the elevated expression of *Alp1* transcripts in colonocytes cultured juxtaposed to epithelium (Supplementary Figure 3D) that complements the SEM analysis (Figure 2B).

### Colon Intestine-Chip supports the multi-lineage differentiation of biopsy-derived colonoids to epithelial monolayer

As the application of organoids in biomedical research is rapidly expanding, several protocols have been developed to differentiate the intestinal organoid-derived cells towards mature cells of the absorptive and secretory lineage. These methods are based either on the engraftment of the organoids in animal models^11^, or on the support of signaling pathways required to define the cell fate in the “crypt-villus” axis. Among the latter, the main approach involves activation of Wnt and inhibition of p38 pathways^26^, often also coupled with inhibition of the Notch pathway^27^.

In the current study, all the stemness factors were maintained in the culture medium over the length of the study. The epithelial culture was maintained in the WENR medium. Immunofluorescence on Day 8 of culture depicts physiological differentiation of the colonoid-derived epithelial cells towards absorptive enterocytes, goblet, and enteroendocrine cells (Figure 3A, Supplementary Figure 2A). This finding was verified by qPCR for characteristic markers of the absorptive and secretory epithelial cell subsets (Figure 3C, Supplementary Figure 2C). As can be deduced, the multilineage differentiation occurred at the expense of proliferating cells, as expected from the life cycle of these cells. Reduced expression of the cycling Intestinal Stem Cells (ISCs) marker, Lgr5, over the course of the culture (Figure 3C, Supplementary Figure 2C), and drastic elimination of EdU positive cells by Day 5 of culture (Figure 3B, Supplementary Figure 3B) are in further support of the gradual depletion of progenitor cells. The presence of endothelium promoted the acquisition of a mature epithelial phenotype (Supplementary Figure 3D). Importantly, the Colon Intestine-Chip established by three different healthy individuals, represented the expected donor-to-donor variability in the differentiation process. This was evident on the varying representation of the absorptive and secretory lineage among the three donors at the completion of the culture. Notably, donors that presented with higher proliferative capacity had lower differentiation towards the absorptive epithelial line and vice versa (Supplementary Figure 2A, B, C).

**Figure 3.**
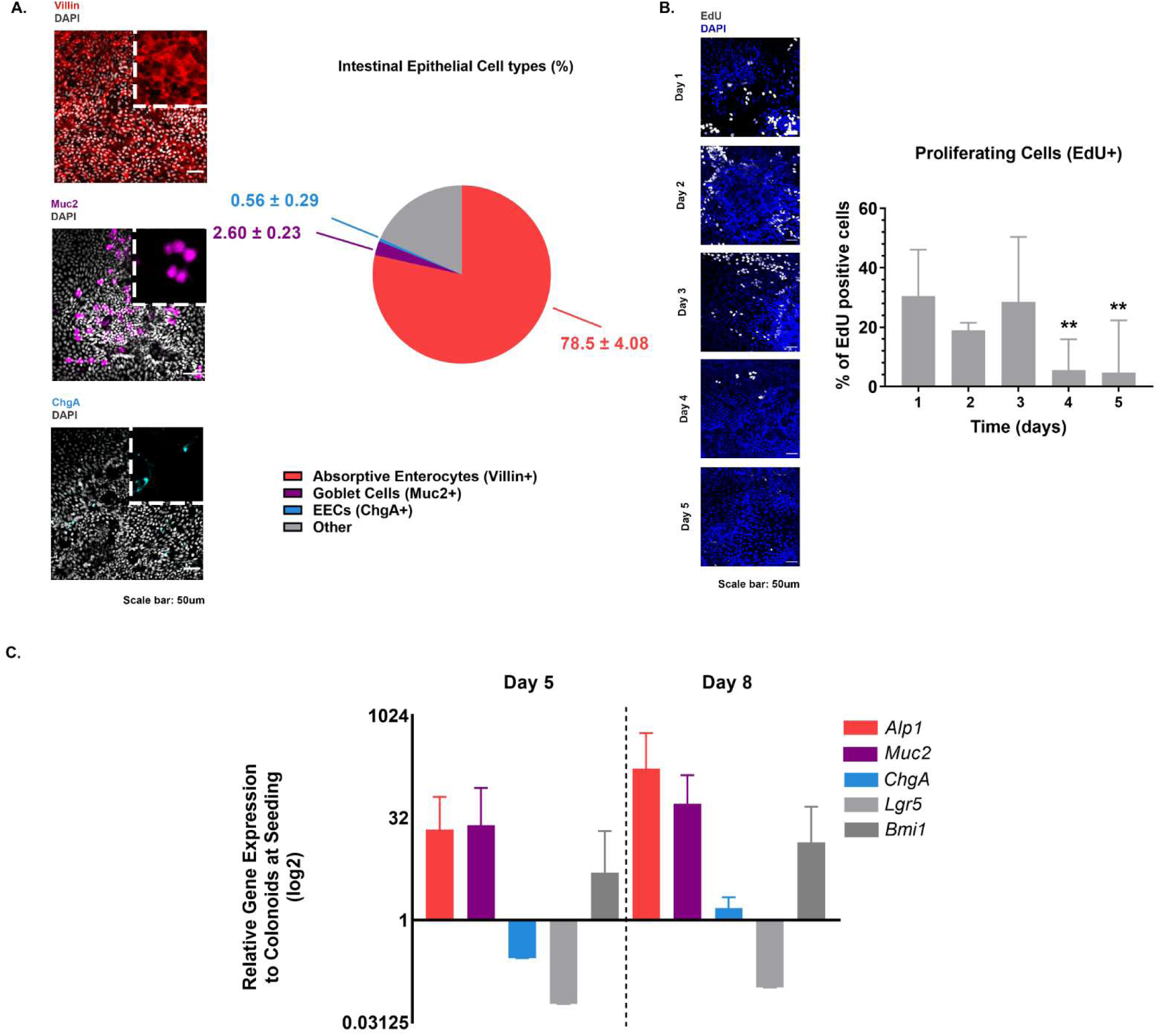
On-Chip culture supports the multi-lineage differentiation of the colonic epithelial cells. **[A]** Representative confocal fluorescent images showing the presence of all the major colonic epithelial cell types on Day 8 of the Colon Intestine-Chip culture. Absorptive enterocytes are stained with anti-Villin (red), Goblet cells with anti-Muc2 (magenta) and enteroendocrine cells with anti-Chromogranin A (cyan). Cells nuclei are shown in gray. Scale bar: 50 μm. The pie chart plots the percentage of the respective cell types. The total number of cells is evaluated based on the DAPI staining. The quantification was performed across 5 different fields of view in 3 individual chips for each cell type. Values are presented as mean±SD. **[B]** Representative confocal fluorescent images indicating the abundance of proliferating cells over the course of the first 5 days of culture. Proliferating cells are stained with EdU (gray) and cells nuclei are shown in blue. Scale bar: 50 μm. The total number of cells is evaluated based on the DAPI staining. The quantification was performed across 3 different fields of view in 3 individual chips for each day. Percentages are plotted as mean±95% CI. The abundance of the proliferating cells is significantly decreased on Day 4 and 5 of the culture. One-way ANOVA, Tukey’s post hoc test, **: p<0.01. **[C]** Identification of each epithelial cell type by qPCR for the Alkaline Phosphatase, (*Alk1*), Mucin 2 (*Muc2*), Chromogranin A (*ChgA*), Leukine-rich-repeat-containing G-protein coupled receptor 5 (*Lgr5*) and polycomb complex protein BMI-1 (*Bmi1*) genes respectively. The relative gene expression values (RGE) of each gene at the respective timepoints (Day 5 and Day 8) are normalized to colonoids at seeding. 18S has been acquired as the house keeping gene. Values are plotted as mean±95% CI on a log2 scale. The expression of indicative for the mature enterocytes and quiescent stem cells genes increase over the course of the culture whereas the expression of the cycling stem cells driving gene Lgr5 decreases.

### Establishment of the barrier disruption model in the Colon Intestine-Chip

The intestinal epithelial barrier maintains mucosal homeostasis via compartmentalization of the resident commensal bacteria from the host immune cells. Activation of the immune cells following disruption of the epithelial barrier results in cytokine release that propagate the epithelial injury. IFNγ is a type II interferon that is produced by immune cells and is increased in inflammatory diseases^28^. IFNγ is produced in the intestinal lamina propria by various cells including Th1 cells and has been implicated in the pathogenesis of IBD^28, 29^. Early studies have revealed the ability of IFNγ to target the intestinal epithelial barrier function by decreasing the transepithelial resistance of T84 monolayers in vitro^9^. Here we administered different concentrations of IFNγ in the basolateral side of the epithelial monolayer, via flow through the bottom channel of the Colon Intestine-Chip, to simulate the *in vivo* action of this cytokine. Fourty eight hours post stimulation, we observed a concentration-dependent increase in the epithelial apparent permeability to 3 kDa Dextran (Figure 4B, Supplementary Figure 3B), together with degeneration of the morphology of the epithelial cells (Figure 4A). Fluorescent staining for F-actin identified compromised cytoarchitecture, depicted by poorly defined cell borders and an increase of enlarged, squamous shaped cells (Figure 4C), whereas immunofluorescence staining revealed displaced localization of ZO-1 and Claudin 4. E-cadherin and Occludin staining in TJs was reduced, in contrast to its increased cytoplasmic expression. This finding is reminiscent of the effect of co-treatment of T84 cells with IFNγ and TNFα, where reduced affiliation of Occludin, JAM-1 and E-cadherin with membrane lipid rafts triggers redistribution of tight junctions proteins and weakens the cell-cell adhesion^29^ (Figure 4C). Further, in accordance to previous publications^29^, basolateral challenge of the IECs with IFNγ resulted in increased Caspase 3 cleavage and activity (Figure 4D, Supplementary Figure 3C). In support of the latter, prolonged exposure to IFNγ resulted in decreased abundance of epithelial cells across the chip surface (Supplementary Figure 3A). Furthemore, we detected elevated secretion of IL-6 24 h post stimulation with IFNγ, and increased levels of serum amyloid A (SAA)^30^ in the effluent of the vascular channel. This finding delineates the propagation of the inflammatory response triggered by the injured barrier. Interestingly, we detected Intercellular Cell Adhesion Molecule-1 (ICAM-1)^31^ in the effluent of the top channel (Figure 4E), described in several cell lines and in the sera of patients in inflamed conditions^32^.

**Figure 4.**
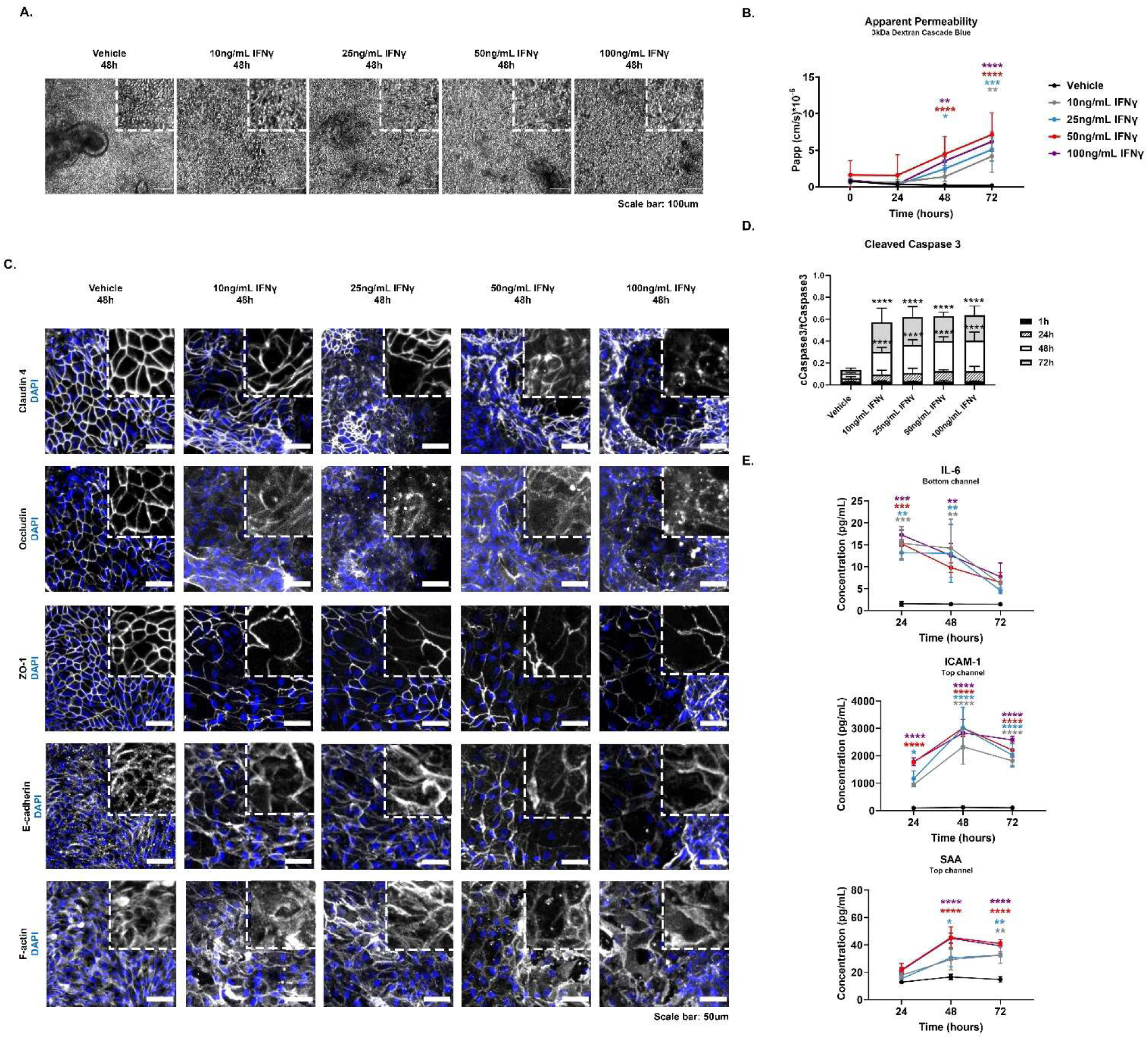
The established epithelial barrier responds on a time and concentration dependent manner to IFNγ, a barrier disruptive agent. **[A]** Representative phase contrast images of the colonic epithelial monolayer 48 h post-treatment with IFNγ. A gradual degeneration of the morphology of the epithelial cells is observed as the concentration of IFNγ increases from 10 ng/mL to 100 ng/mL. Scale bar: 100 μm **[B]** Apparent Permeability (P_app_) of the colonic epithelial cells to 3 kDa Dextran Cascade blue, over the course of 72 h of basolateral stimulation with IFNγ. A dose and time dependent disruption of the barrier integrity is observed. Data shown correspond to one representative out of three independent experiments. n= 3 - 5 Chips/condition, mean±95% CI, Two-way ANOVA, Tukey’s post hoc test, **: p<0.01, ****: p<0.0001. Scale bar: 50 μm. **[C]** Representative confocal immunofluorescence images indicating the dose dependent degeneration of the epithelial tight junctions, as shown by the redistribution of Claudin 4, ZO-1 and F-actin, and internalization of Occludin and E-cadherin. Cells nuclei are shown in blue. **[D]** Quantification of the cleaved over the total Caspase 3 in cell lysates from the epithelial channel of the Colon Intestine-Chip. A significant and concentration dependent increase in the levels of cleaved Caspase 3 is observed 48 h and 72 h post stimulation. n= 3 Chips/ condition mean±95% CI, Two-way ANOVA, Tukey’s post hoc test, ****: p<0.0001. **[E]** Time course secretion of cytokines and vascular injury related serum proteins in the Colon Intestine-Chip. A concentration and time dependent secretion of IL-6, and ICAM-1 and SAA, in the bottom and top channel respectively, is observed upon treatment with IFNγ. Culture effluents were collected at different time intervals. n= 3 Chips/ condition mean±SD, Two-way ANOVA, Tukey’s post hoc test, *: p<0.05, **: p<0.01, ***: p<0.001, ****: p<0.0001.

### IL-22 signaling is preserved in the Colon Intestine-Chip

IL-22, an immune cell derived cytokine, has been implicated in both protection and worsening of mouse models of colitis depending on disease models^33, 34^. Thus a better understanding of IL-22 in human systems may be informative. As previously reported, IL-22 binds to its receptor at the basolateral side of polarized IECs^35^ and triggers the phosphorylation of STAT3^36, 37^. We first confirmed gene expression of the two IL-22 receptor subunits, *Il22ra1* and *Il10rb*, based on RNA-Seq data from organoid-derived epithelial cells cultured in the presence or absence of endothelium and/or stretch (Figure 5E). Interestingly, the expression of the IL-22 receptor was higher in the organoids cultured on the chip, as compared to those in suspension culture. The expression of both *IL22ra1* and *IL10rb* in the organoid-derived epithelial cells in the Colon Intestine-Chip, was also higher in the presence of the endothelium. This finding is in agreement with all our experiments that signify the supportive effects of endothelium. These data highlight the potentially incomplete understanding of human IL-22 biology based on data derived from organoids cultured in suspension^22^.

**Figure 5.**
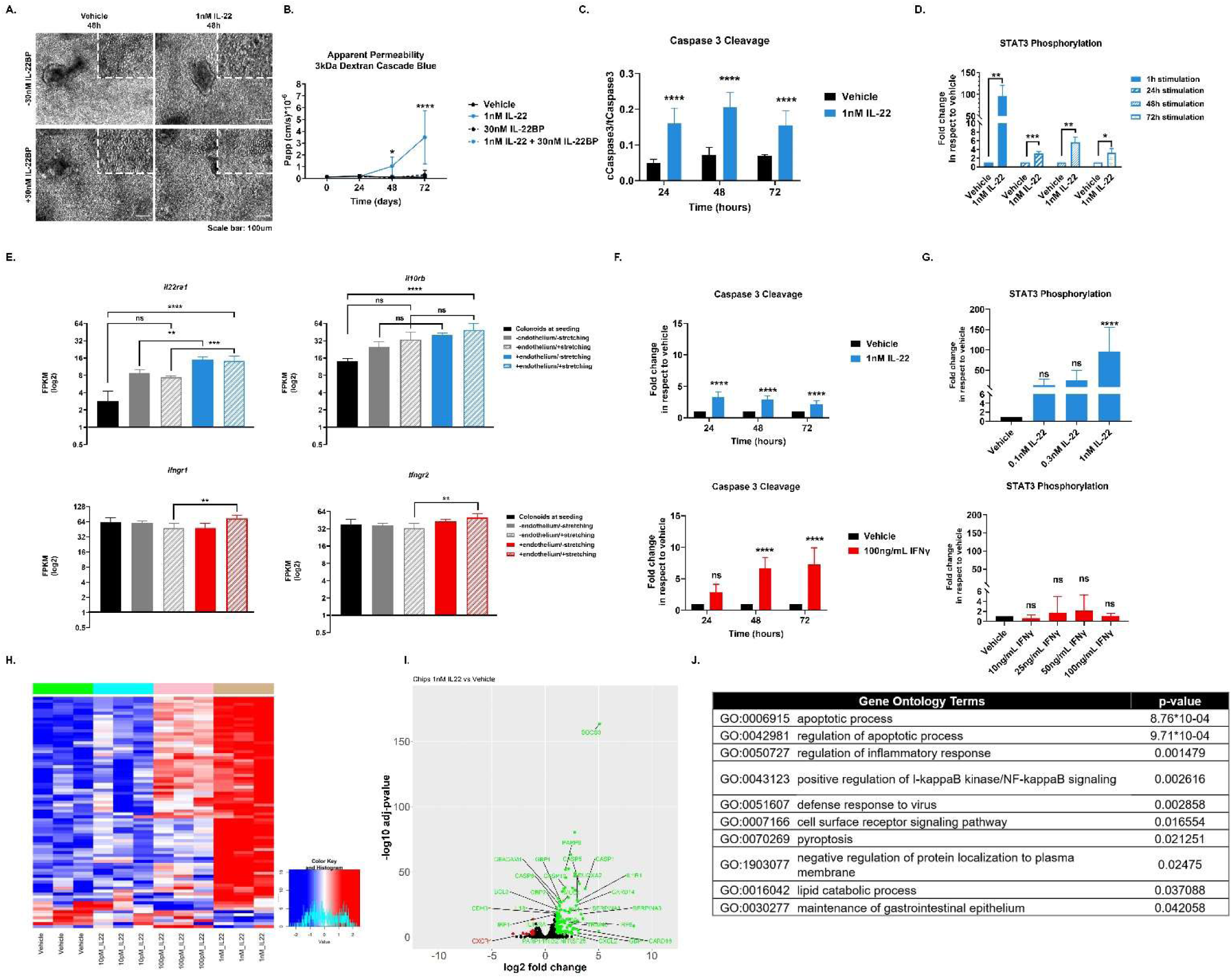
IL-22 induces disruption of the epithelial barrier in the Colon Intestine-Chip, an effect rescued by the presence of its soluble receptor IL-22BP. **[A]** Representative phase contrast images of the colonic epithelial monolayer 72 h post treatment with 1 nM IL-22. The basolateral stimulation of the Colon Intestine-Chip results in compromised morphology of the epithelial cells. The epithelial cells morphology is rescued in the presence of 30 nM IL-22BP. Scale bar: 100 μm. **[B]** Apparent Permeability (P_app_) of the colonic epithelial cells to 3 kDa Dextran Cascade blue, over the course of 72hrs basolateral stimulation with 1 nM IL-22 and/ or 30 nM IL-22BP. IL-22 triggers loss of the epithelial barrier integrity which is rescued upon co-treatment with IL-22BP. n= 3-6chips/condition, mean±95% CI, Two-way ANOVA, Tukey’s post hoc test, *: p<0.05, ****: p<0.0001. **[C]** Quantification of the cleaved over the total Caspase 3 in cell lysates from the epithelial channel of the Colon Intestine-Chip. A significant and sustained increase in the levels of cleaved Caspase 3 is observed over the course of 72 h basolateral stimulation with 1 nM IL-22. n= 3 Chips/ condition mean±95% CI, Two-way ANOVA, Tukey’s post hoc test, ****: p<0.0001. **[D]** Quantification of the phosphorylated over the total STAT3 in cell lysates from the epithelial channel of the Colon Intestine-Chip, expressed as fold increase over the Vehicle, reveals a sustained activation of STAT3 over the course of 72 h basolateral stimulation with 1 nM IL-22. n= 3 Chips/ condition mean±95% CI, Unpaired t-test performed for each timepoint, *: p<0.05, **: p<0.01, ***: p<0.001. **[E]** The gene expression of the two subunits of the transmembrane IL-22 receptor, *Il22ra1* and *Il10rb*, and the transmembrane IFNγ receptor, *Ifngr1* and *Ifngr2*, was confirmed in the epithelial channel on Day 5 of culture of the Colon Intestine-Chip. The on-chip culture enhances the expression of both receptors in the colonic epithelial cells. The graph represents FPKM values generated from bulk RNAseq analysis of the epithelial cells and plotted on a log2 scale. n= 3-7 Chips/ condition, mean±95% CI, One-way ANOVA, Tukey’s post hoc test, ns: p>0.05, **: p<0.01, ***: p<0.001, ****: p<0.0001. **[F]** Quantification of the cleaved over the total Caspase 3 in cell lysates from the epithelial channel of the Colon Intestine-Chip, expressed as fold increase over the Vehicle, reveals a cytokine and time dependent activation of apoptosis over the course of 72 h basolateral stimulation with either 1 nM IL-22 or 100 ng/mL IFNγ. n= 3-6 Chips/ condition mean±95% CI, Two-way ANOVA, Tukey’s post hoc test, ns: p>0.05, ****: p<0.0001. **[G]** Quantification of the phosphorylated over the total STAT3 in cell lysates from the epithelial channel of the Colon Intestine-Chip, expressed as fold increase over the Vehicle, reveals a cytokine specific activation of STAT3 upon basolateral stimulation with various concentrations of IL-22 and IFNγ. n= 3 Chips/ condition mean±95% CI, One-way ANOVA, Tukey’s post hoc test, ns: p>0.05, ****: p<0.0001. **[H]** Heat map of the 1000 most variant genes. Samples cultured in the presence of various concentrations of IL-22 are clustered separately from vehicle condition. n=3 Chips/condition. **[I]** Volcano plot of the differentially expressed genes comparing epithelial cells on day 5 of culture under baseline conditions and upon basolateral stimulation with 1nM IL-22. Genes upregulated and downregulated in the presence of endothelium are shown as green and red respectively. **[J]** Gene Ontology biological processes analysis, comparing epithelial cells on day 5 of culture treated with 1 nM IL-22 or vehicle, indicates the upregulation of terms related to the inflammatory response and apoptotic pathway.

Next, we stimulated, the epithelial cells in the Colon Intestine-Chip on Day 5 of culture, through the basolateral side with either 0.1 or 1 nM of IL-22 in the presence or absence of the IL-22 soluble binding protein IL-22BP, for 1 h. A significant increase in the phosphorylation of STAT3 was observed upon challenge with IL-22 at the highest concentration used (1 nM), an effect abolished by co-culture with IL-22BP (Data not shown). Maximal activation of STAT3 was obtained in the acute phase of the treatment (1 h). STAT3 activity was still evident upon prolonged stimulation (up to 72 h) (Figure 5D), indicating the sustained sensitivity and responsiveness of the system for potential secondary readouts. Unlike the downstream effects of IL-22, treatment with IFNγ did not result in significant activation of STAT3, highlighting the specificity of the event (Figure 5G).

### IL-22 acts as a barrier disruptive agent for the mature colonic epithelial monolayer in the Colon Intestine-Chip

IL-22 is regulated by IL-12 and IL-23, and is central to the pathogenesis of IBD. The exact role and directionality of the IL-22 effects in intestinal epithelial homeostasis is not clearly defined. Current reports describe anti-inflammatory effects, induction of homeostatic factors or epithelial cell proliferation^37^, as well as reduction in epithelial integrity either in mice models or the Caco2 cell line^21, 23^. A recent study showed, interestingly, induction of endoplasmic reticulum (ER) stress response in healthy and IBD colonoid-derived epithelial cells^22^. We next used the Colon Intestine-Chip platform to assess the effects of IL-22 on the epithelial barrier, as described above with the reproducible effects of IFNγ. As shown (Figure 5B), challenge with IL-22 (1 nM) for 48 h (1 nM), resulted in increased apparent permeability of the colonic epithelium to 3 kDa Dextran. IL-22 (1 nM) induced permeability was abrogated when cotreated with IL-22BP (30 nM) (Figure 3B). We found compromised morphology of the epithelial monolayer (Figure 5A), with poorly defined cell borders and increase of floating cells in the effluent. Finally, we show that IL-22 increases cleavage of Caspase 3 (Figure 5C), affecting apoptosis similar to IFNγ (Figure 5F). These data were supported by transcriptomic analysis and downstream functional characterization of the differentially regulated genes (Figure 5I). Functional enrichment of GO terms, indicate upregulation of biological processes associated with apoptosis and inflammatory responses (Figure 5H, I, J). Importantly, our data strongly suggest that IL-22 targets the human colonic epithelial cells and disrupts the barrier function, acting as an injurious, rather than a homeostatic or anti-inflammatory factor^37–40^. Notably, the IL-22 driven effects obtained with the Colon Intestine-Chip refer to a healthy state. Investigation of the potential compensatory actions of IL-22 in the inflamed epithelium is beyond the scope of this study.

## DISCUSSION

Gastrointestinal diseases affect all ages and account for a significant part of acute and chronic diseases across the world, with more than 11% of the US population estimated to suffer from a digestive disorder. A number of these diseases compromise the quality of life and often are associated with decreased life expectancy. Although advancements in population screening and the diagnostic methods have improved the overall prognosis^41^, a number of these diseases remain as unmet medical needs.

Intestinal epithelial homeostasis is intimately linked to the development of a tight epithelial barrier, owed to the specialized TJs connecting adjacent polarized epithelial cells and associated supporting factors, such as the mucin layer. The significance of the barrier function is clearly demonstrated by the devastating consequences from its disruption, including the development of inflammatory diseases such as IBD. Our understanding on the gastrointestinal physiology and disease processes has dramatically increased lately, but the translation of these findings to efficient therapeutics and associated biomarkers still lags behind. A significant reason for the latter has been the limited availability of human cell systems that recapitulate cell interactions and functionality of the human tissue of origin. The recent development of organoids, as patient-specific cell sources with unrestricted potential for propagation, has revolutionized our capabilities for disease modeling. A major advantage of organoids is the relatively fast potential to uncover the spectrum of disease phenotypes by direct comparisons between patients’ samples. A caveat of the system is that they lack the ability to maintain the tissue microenvironment and recapitulate the cytoarchitecture, significant for cell-cell interactions and identification of disease mechanisms for successful targeting^42^.

One of the most developed organoid systems is the gut, owing to its unique ability for life-long renewal from the villous crypts, enables the reproducible establishment of organoids from the small and the large intestine^43, 44^. An important restriction of the organoids is that they do not allow for direct access to the apical side of the cells, which faces inward of the enclosed cystic cell structure^44^. This becomes a bottleneck, in particular with the IECs, as the strictly polarized expression of key proteins for development, homeostatic or disease-specific functions, is linked to disease-mediating mechanisms. To address this need, methods for expansion of 3D organoids into monolayered epithelial structures were established in the transwell culture systems^20, 45–47^. These developments enabled studies on infectious agents, a major and emergent group of the GI diseases, and co-culture of different cell populations, although in a non - dynamic microenvironment. In the current study, we sought to address this need by leveraging the developments in microphysiological, Organ-on-Chip systems and the human organoids technology to develop a Colon Intestine-Chip, as a platform for preclinical studies. We have previously reported in a microengineered Duodenum Intestine-chip, the method for establishing a juxtaposed culture of human organoid-derived epithelial monolayer and primary intestinal mircovascular endothelial cells on tissue relevant-derived extracellular matrix. The microenvironment of the Colon Intestine-Chip include exposure to shear stress and stretching to recapitulate major aspects of the intestinal peristaltic motion. Although, unlike other protocols described previously^44, 46, 48^, none of the stemness supporting factors are removed during the course of the culture, the organoid-derived epithelial cells transition from a proliferative to a fully differentiated state. Further, the presence of endothelium from the beginning of the culture was found to support the earlier establishment of a tight epithelial barrier, a consistent finding in all donors tested. As described in the results, upregulation of key signaling pathways related to epithelial differentiation, metabolism, and ion transportation, unveil the physiological relevance of the transcriptomic phenotype in the Colon Intestine-Chip. The robustness and reproducibility of the protocol, and the demonstrated inter-individual variability in the pace of differentiation of colonoids to epithelial cell subtypes were highly suggestive of the potential of the Colon Intestine-Chip as a platform for human studies. For further confirmation of the latter, we proceeded with assessment of the functional response of the Colon Intestine-Chip to a well-established barrier disruption cytokine, IFNγ^9^. As has been shown by a number of studies with cell lines and *in vivo* experimental models, IFNγ modulates the integrity of both the intestinal epithelial^9^ and endothelial^49^ barrier. The disruption of the epithelial barrier, specifically, is driven by both apoptosis^29^ and cytoskeletal remodeling, via upregulation of the MLCK^8, 9^. We found that upon exposure to IFNγ, the Colon Intestine-Chip showed loss of the epithelial barrier integrity in a concentration- and time-dependent manner, a response reproduced in all donors employed in this study. Activation of apoptosis, as indicated by the cleavage of Caspase 3, and erosion of the epithelial tight junctions were detected as early as 48 h post stimulation in comparison to previously described models using T84 and Caco2 monolayers^8,9,29^. These results indicate the response and sensitivity of the Colon Intestine-Chip platform to a well-established, disease relevant, barrier disruption bioassay and provide reassurance for its potential to complement and further advance existing models.

Next, we applied the Colon Intestine-Chip to elucidate the effect of IL-22, a member of the IL-10 family of cytokines secreted in humans primarily by subpopulations of CD4+ T cells, including Th22, Th17, Th1 and γδT cells, as well as ILC3s, NK and dendritic cells (DCs)^34, 50–55^. Elevated levels of IL-22 have been detected in the intestinal mucosa of IBD and in the serum of CD patients^56^. IL-22 binds to its dimeric transmembrane receptor, consisting of the IL-10Rb and IL-22Ra1 subunits^57^ and activates downstream signaling resulting in the phosphorylation of STAT3.

Although a number of reports advocated for a protective role of IL-22 in the inflammatory state, as it promotes cell survival and proliferation as well as secretion of mucins, chemokines, and antimicrobial peptides of the S100 and REG families^58–61^, the overall effects of IL-22 in the epithelial barrier homeostasis, as per animal and human studies remain unclear^62, 63^. Our findings in the Colon Intestine-Chip showed concentration-dependent increase of epithelial permeability in response to IL-22, consistent with studies in transwells with Caco2 cells^21^, while they do not support the previously described regenerative effect of IL-22 on the crypt stem cells^39^. IL-22 acts via activation of STAT3 and STAT1. It has been hypothesized that the cross talk between type I IFNs and the members of the IL-10 cytokine family, IL-10 and IL-22, may alter the downstream pathway landscape by enhancing a STAT1 mediated proinflammatory response^64–66^. Moreover, recent reports indicate an IL-22 elicited inhibition of Wnt and Notch pathway in small intestinal organoids, that limits the renewal of the Lgr5^+^ stem cell pool^36^. These data collectively suggest a biological action of IL-22 dependent upon the specific characteristics of the employed experimental model and further support the detrimental effect of IL-22 during a time-window, where little self-renewing cells are still present in the Colon Intestine-Chip. Independent of its short-term effects, prolonged exposure to IL-22 has been associated to chronic diseases, such as intestinal inflammation and cancer^67, 68^. Additional complexity is added by secretion from a subset of DCs in both human and mouse^69^, of a soluble IL-22 binding protein (IL-22BP) which binds IL-22 with higher affinity than transmembrane receptor^55^. Blockade of the IL-22 effect on STAT3 activation, by co-treatment of the Colon Intestine-Chip with its soluble binding protein, IL-22BP, shows the specificity of the effect and suggests that this platform can be employed to target tissue-specific transcription factors.

In summary, we report here the development of a Colon Intestine-Chip, a microphysiological system that provides the microenvironment to enable differentiation of human organoids to an epithelial monolayer of polarized cells forming a tight barrier. We show that this human platform can be used to study potential barrier disruptive factors in a reproducible and specific manner. The implications of these findings for understanding the regulation of barrier function are very important given the list of conditions causing leaky barrier and the association of the latter with degenerative and other diseases.

## METHODS

### Cell Culture

Human biopsy derived colonoids, from two male and one female donor, isolated according to experimental protocols approved by Johns Hopkins University Institutional Review Board (IRB# NA_00038329) were provided by Prof. Mark Donowitz’ s group.

Colonoids were cultured in Matrigel (Corning, 356231) and in the presence of the commercially available medium IntestiCult™ Organoid Growth Medium (Human) (STEMCELL Technologies, 06010), supplemented with the antibiotic primocin (Invivogen, ant-pm-2) in a concentration of 1: 500 (vol/ vol). Passaging was performed every 7 days, colonoids were recovered from the Matrigel, using Cell Recovery Solution (Corning, 354253), and fragmented for 2 min at 37°C in a digestion solution containing TrypLE (Thermo Fischer Scientific, 12604013), diluted with PBS1X 1:1 (vol/vol), and supplemented with 10 μM Y-27632). 10 μM Y-27632 (Sigma-Aldrich, Y0503) and 5 μM CHIR99021 (Stemgent, 04-0004-10) were present in the culture medium for 72 h post passaging.

Colonic Human Intestinal Microvascular Endothelial Cells (cHIMECs) were obtained by Cell Biologics (H-6203) and Cell Systems (ACBRI 666). cHIMECs were cultured in gelatin (Cell Biologics, 6950) coated flasks in Endothelial Cell Growth Medium MV2 (EGM-MV2) (Promo Cell, C-22121), which contained 5 ng/mL recombinant human epidermal growth factor (rhEGF), 10 ng/mL recombinant human basic fibroblast growth factor (rhbFGF), 20 ng/mL long R3 insulin-like growth factor (R3-IGF), 0.5 ng/mL recombinant human vascular endothelial growth factor 165 (rhVEGF), 1 μg/mL ascorbic acid, 0.2 μg/mL hydrocortisone, 5% fetal bovine serum (Sigma Aldrich, F4135) and 1:1,000 (vol/ vol) primocin (Invivogen, ant-pm-2).

Cells were cultured in a humidified environment, at 37°C in 5% CO_2_. For the experimentation, human colonoids of passage 30 or below, and cHIMECs of passage 7 or below, were used.

### Establishment of the Colon Intestine-Chip

The Colon Intestine-Chips (Emulate Inc., 10231-2), fabricated by Emulate Inc., consisted of two parallel channels separated by a porous PDMS membrane (diameter: 7 μm, spacing: 40 μm), and two lateral channels where vacuum is applied. The dimensions of the top channel, where colonic epithelial cells reside, are 1000 μm x 1000 μm (width x height), its volume is 28.041 μL and its cell culture surface of 28 mm^2^. The endothelial cells were introduced in the bottom channel the dimensions of which is 1000 μm x 200 μm (width x height), its volume equal to 5.584 μL and its cell culture surface equal to 24.5 mm^2^. The interface of the two cell types extended in an area of 17.1 mm^2^. The timeline of the establishment of the Colon Intestine-Chips was the following (Fig. 1):

*Day -1*: The PDMS membrane was activated using ER1 and ER2 reagents (Emulate, 10465). A solution of 0.5 mg/mL ER1 in ER2 was prepared and 50 μL was introduced in each channel. Chips were incubated under UV light for 10 min, followed by 3 min of incubation in room temperature and another 10 min of UV treatment. Subsequently, both channels were washed with ER2, then PBS 1X, and coated overnight at 37°C, with 200 μg/mL Collagen IV (Sigma Aldrich, C5533) and 100 μg/mL Matrigel (Corning, 356231), and 200 μg/mL Collagen IV (Sigma Aldrich, C5533) and 30 μg/mL Fibronectin (Corning, 356008), in the top and bottom channel respectively.

*Day 0*: In order to be seeded on the Chips, colonoids were recovered from Matrigel using Cell Recovery Solution (Corning, 354253), and fragmented by incubation at 37°C for 2 min, in a digestion solution containing TrypLE (Thermo Fischer Scientific, 12604013), diluted with PBS1X 1:1 (vol/vol) and supplemented with 10 μM Y-27632). They were resuspended in IntestiCult™ medium, supplemented with 10 μm Y-27632 (Sigma-Aldrich, Y0503) and 5 μm CHIR99021 (Stemgent, cat# 04-0004-10) and introduced in the apical channel of the Colon-Chip, in a seeding density of 7 x 10^6^ cells/ mL. Y-27632 (10 μM) and CHIR99021 (5 μM) are maintained in the culture medium for 72 h thereafter.

When investigating the interaction between epithelial and endothelial cells, cHIMECs were seeded on the basal side of the PDMS membrane prior to the seeding of epithelial cells in the top channel. Briefly, 20μL of the suspension of endothelial cells, in a seeding density of 8×10^6^ cells/mL were introduced in the bottom channel and Colon Intestine-Chips were incubated inverted at 37°C, in humidified environment for up to 1 h. The bottom channel was washed and thereafter immersed in EGM-MV2 medium.

*Day 1*: Both channels were washed with the corresponding media in order to remove the non-adherent cells. Colon Intestine-Chips were attached to devices called Pod™ Portable Modules (Emulate Inc., 10153) and connected in the automated cell culture system Zoë™ Culture Module (Emulate Inc.). Each Zoë is connected to an Orb^TM^ Hub Module that supplies the gas, vacuum, and power. The flow rate was set at 60 μL/h in both channels.

*Day 2*: Medium without Y-27632 and CHIR99021, was replenished in both channels.

*Day 3*: Colon Intestine-Chips were acclimated to mechanical forces, using 2% strain, 0.15 Hz frequency.

*Day 4*: Stretching was increased to 10% strain, 0.15 Hz frequency, and culture medium was replenished. Both channels were perfused at 1000 μL/h for 5 min to flush out any detached cells.

*Day 5-8*: Culture media was refreshed, and detached cells were removed by flushing (1000 µL/h, 5 min) every other day of the culture.

### Assessment of Apparent Permeability

The epithelial barrier integrity was assessed based on the apparent permeability of 3kDa Dextran Cascade Blue molecule (ThermoFischer, D7132). Dextran Cascade Blue was added in the epithelial medium (apical channel) in a concentration of 100 μg/mL. The concentration of the dye, that had diffused in the bottom channel, was monitored, based on a standard curve and the absorbance at of the bottom channel effluents at 375-420 nm. For the calculation of the apparent permeability (P_app_) the following mathematical formula was used:

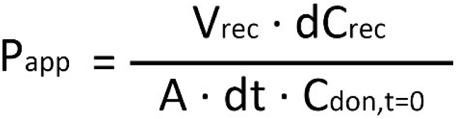

Where P_app_ is the apparent permeability (cm/s), V_rec_ is the volume of the retrieved effluent from the vascular channel (mL), dC_rec_ is the difference of the concentration of the dye in the effluent and input medium of the vascular channel (mg/mL), A is the surface areaon which the diffusion of the dye occurs (cm^2^), dt is the time period on which the P_app_ is assessed (s), and C_don, t=0is_ the concentration of the dye in the dosing epithelial medium (mg/mL).

### Cytokines Stimulation

IL-22 (R&D, 782-IL-010) in the presence or absence of 30 nM IL-22BP (R&D, 8498-BP-025) and IFNγ (Peprotech, 300-02) were introduced in the culture medium of the endothelial channel at various concentrations. The duration of the treatment differed based on the readout analysis. The appropriate vehicle controls were used in each stimulation experiment.

### Immunofluorescence

At the indicated timepoints Colon Intestine-Chips were fixed at room temperature, for 15 min in 4% Paraformaldehyde (PFA) solution, or at -20°C for 20 min in ice-cold methanol, according to the target epitope. Briefly, chips were washed in both channels with PBS1X, immersed in blocking solution (10% Normal Donkey Serum, 0.1% Triton X-100 in PBS1X) for 15-30 min at room temperature, and incubated with primary antibody, overnight at 4°C. Subsequently, chips were stained with the appropriate combination of secondary Alexa Fluor antibodies (Abcam) and counterstained with 50ug/ml DAPI (Invitrogen, D1306). The antibody solutions were prepared using 5% Normal Donkey Serum in PBS1X, in the dilution ratios mentioned in Supplementary Table 1. In case of the morphological analysis of the epithelial cytoarchitecture, F-actin was stained with Phalloidin, conjugated with the appropriate Alexa Fluorophores (Abcam), which were added in the secondary antibody solution. To assess epithelial cell proliferative capacity, chips were treated with EdU (20 µM), and cells were harvested and assessed 24 hrs later per manufacturing instruction (ThermoFischer, cat#C10337).

### RNA isolation, Reverse Transcription, and qPCR

Each channel of the Colon Intestine-Chip was lysed using TRI Reagent® (Sigma-Aldrich, T9424) and RNA was isolated using the Direct-zol RNA Purification Kit (Zymo Research, R2060). Genomic DNA was removed using the TURBO DNA-free™ Kit (ThermoFischer, AM1907) and reverse transcription to cDNA was performed with the SuperScript IV Synthesis System (ThermoFischer, 18091050). TaqMan Fast Advanced Master Mix (Applied Biosystems, 4444963) and TaqMan Gene Expression Assays (Supplementary Table 2) were used for the Quantitative Real Time PCR (qPCR). The qPCR was detected on a QuantStudio 3 PCR System (Fisher Scientific). The raw data was transformed to linear data by calculating the 2^-ΔΔCt^ values.

### Imaging and Imaging Quantification

Phase contrast images were acquired using the Zeiss XIO vert.A1 brightfield microscope. Fluorescent z-stack images and fluorescent tile images of the Colon Intestine-Chip were acquired using the inverted laser-scanning confocal microscope Zeiss LSM 880 and the epifluorescence inverted microscope Olympus IX83, respectively. Image processing and quantification was performed using Fiji (Version 2.0), as per standard methods.

### Scanning Electron Microscopy

At the indicated timepoints, the Colon Intestine-Chips were fixed at room temperature, for 2 h in 2.5% Glutaraldehyde solution and washed three times with 0.1 M sodium cacodylate (NaC) buffer. Concomitantly, the Chip was trimmed using a razor so that the lateral and top portions of PDMS were removed and the top channel was revealed. Afterwards, the samples were fixed with 1% osmium tetroxide (OsO_4_) in 0.1 M NaC buffer for 1 h at room temperature and dehydrated in graded ethanol. The Chip samples were dried using the chemical drying agent Hexamethyldisilizane (HMDS), sputter coated with platinum and images were acquired using the Hitachi S-4700 Field Emission Scanning Electron Microscope.

### Assessment of STAT3 Phosphorylation and Caspase 3 Cleavage

Following a 1 h stimulation with IL-22, the Colon Intestine-Chip was disconnected from Pods, endothelial cells were enzymatically removed and subsequently epithelial cells were lysed with a protein lysis buffer (MSD, R60TX-3) containing protease (ThermoFischer, 78425) and phosphatase inhibitors (Sigma-Aldrich, P0044 & P5726). Protein extraction was performed according to standard protocols and the quantification of the total amount of protein/ sample was done by Bradford assay (Thermo Fischer, 1856210). Total and phosphorylated STAT3 and total and cleaved Caspase 3, were detected using the MSD technology (Meso Scale Discovery, tSTAT3-K150SND, pSTAT3- K150DIA, c/t-Caspase 3-K15140D).

### Assessment of ECM coating

ECM proteins were stained using an Alexa Fluor 488 conjugated NHS Ester (ThermoFisher, A20100). Following membrane activation and coating, each channel was pretreated with borate buffer (pH 8.5) at room temperature for 5 min and stained with NHS Ester dye solution in borate buffer (1:500) at room temperature for 25 min in dark. Both channels were gently rinsed with PBS 1X before and images were acquired.

### RNAseq Analysis

Total RNA was extracted from 36 samples using TRI Reagent® (Sigma-Aldrich, T9424). Bulk RNA sequencing was performed in colonoids, in suspension culture, and colonoid derived epithelial cells cultured in the presence or absence of endothelium and/ or stretch on days 5 and 8 of culture, using a minimum of three technical replicates per experimental group (Supplementary Table 3).

The bulk RNA sequencing platform used was the Illumina TruSeq paired-end sequencing with maximum read length 2×100 bp. The sequencing depth was at ∼48M paired end reads/sample. The sequence reads were trimmed using Trimmomatic v.0.36 in order to remove poor quality adapter sequences and nucleotides. Using the STAR (Spliced Transcripts Alignment to a Reference) aligner v.2.5.2b, we mapped the trimmed reads to the *Homo* sapiens reference genome GRCh38 (available on ENSEMBL) and generated the corresponding BAM and featureCounts files. Next, using the v.1.5.2 subread package, we calculated the gene hit counts. It is worth mentioning that only unique reads that fell within exon regions were counted. Genes with very low expressions across samples were filtered-out and the remainder was used to form our dataset which was used for the Differential Gene Expression (DGE) analysis. We performed DGE analysis using the “DESeq2” R package by Bioconductor. For all DGE analyses we applied the following widely accepted strict thresholds: p-value<0.01 and |log_2_FoldChange| > 2.

### Functional enrichment analysis

The genes identified from the DGE analysis were used to perform Gene Ontology (GO) enrichment analysis with the GO biological processes (GO BP). The GO terms enrichment analysis was performed using the GO knowledgebase (Gene Ontology Resource http://geneontology.org/).

### Statistical Analysis

Results are expressed as Mean ± 95% CI or Mean ± SD, as appropriate. The statistical analysis was performed using GraphPad Prism, version 8.3.0 (GraphPad Software Inc.), by applying two-tailed, unpaired t-test, one-way or two-way ANOVA followed by Tukey’s post hoc multiple comparison tests, according to the experimental design. Any p-value lower than 0.05 was considered statically significant.

## ACKNOWLEDGEMENTS

We would like to thank Prof. Mark Donowitz for providing the colonic organoid samples. We also thank Brett Clair for scientific illustrations.

## AUTHOR CONTRIBUTIONS

Design, execution and analysis of the experiments A.A.; Contribution to the experiments R.L., A.D., C.L., G.K., I.M.; Design and analysis of the RNAsequencing studies D.V.M., M.P., R.P., E.S.M., G.A.; Interpretation of the bioinformatic data D.V.M., M.P., E.S.M.; Design and analysis of the IL-22 experiments R.P, A.B., A.A.; Interpretation of the data K.K., C.G., A.A.; Significant contribution in the writing of the manuscript A.A., M.P., E.S.M., B.B., G.H.; Strategy and design of the studies and writing of the manuscript K.K., C.G.; All authors have reviewed the published version of manuscript.

---

**Supplementary Figure 1.**
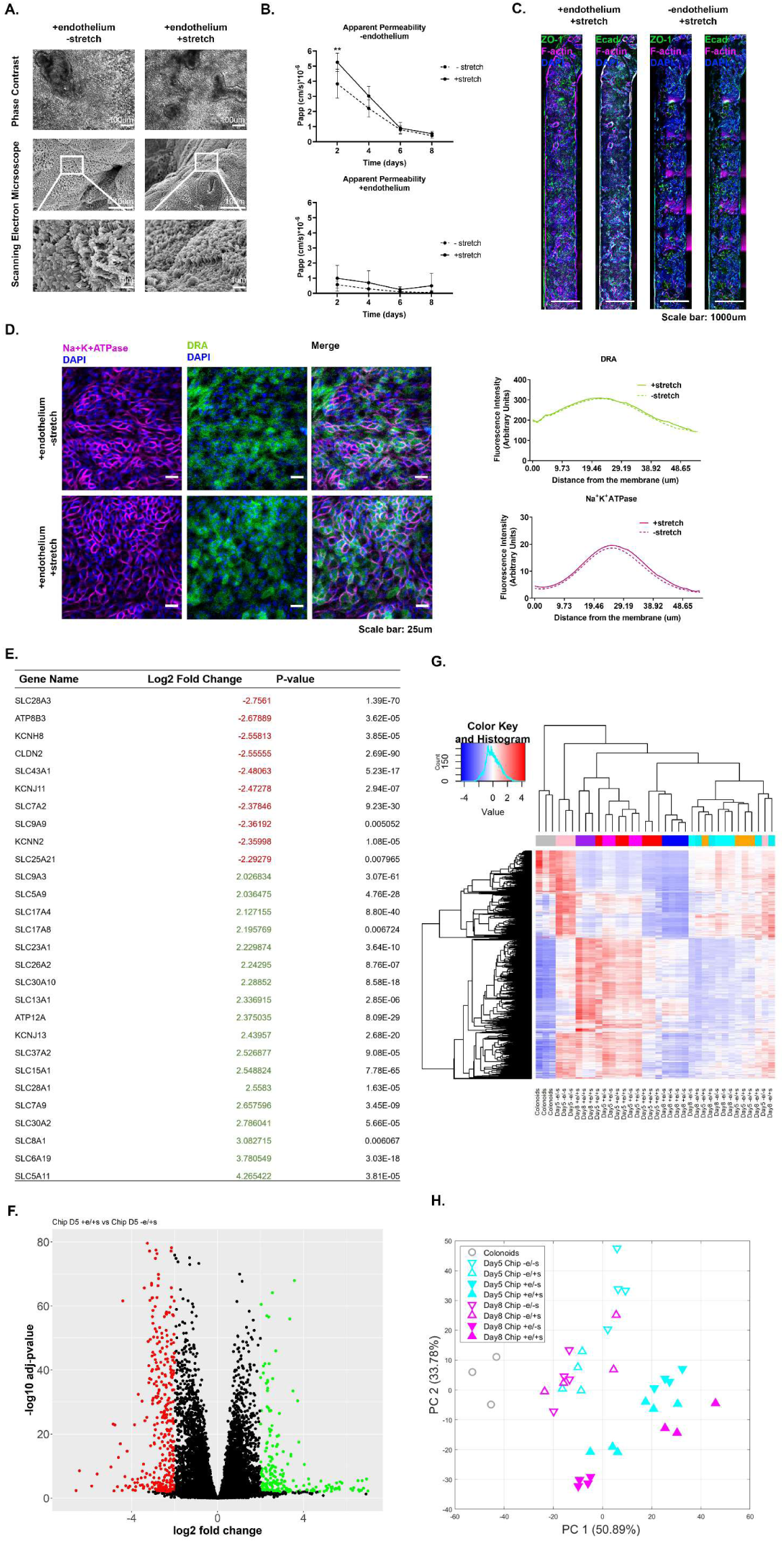
**[A]** Apparent Permeability (P_app_) of the colonic epithelial cells to 3 kDa Dextran Cascade blue, over the course of an 8 day culture, in the presence or absence of stretch (10% strain, 0.15 Hz) and/or endothelium. No significant effect of stretch on the establishment of the epithelial barrier was revealed. Data shown correspond to one representative out of three independent experiments. n= 3-11 Chips/condition, mean±95% CI, Two-way ANOVA, Tukey’s post hoc test, **: p<0.01. **[B]** Representative phase contrast and Scanning Electron Microscopy (SEM) images of the colonic epithelial on day 5 of the Colon Intestine-Chip culture do not reveal a significant contribution of stretching on the maturation of the epithelial brush border in the presence of endothelial cells. **[C]** Representative widefield fluorescent tile images showing the uniform expression of the tight junctions’ proteins, ZO-1 (green) and anti-E-cadherin (magenta), along the Colon Intestine-Chip. Cells nuclei are shown in blue. Scale bar: 1000 μm. **[D]** Representative confocal immunofluorescence images indicating the basolateral localization of the transporter Na^+^K^+^ATPase (magenta) and the apical localization of the transporter DRA (green). Cells nuclei are shown in blue. The application of stretch does not affect the establishment of the polarity of the epithelial cells. The plots indicate the mean fluorescent intensity distribution for each channel, DRA (green) or Na^+^K^+^ATPase (magenta), along the basal-apical axis of the epithelial cells. Three FOVs were acquired per chip. n= 3 chips/ condition. Scale bar: 25 μm. **[E]** Table of the upregulated (green) or downregulated (red) transporter and tight junction proteins genes on epithelial cells on day 5 of culture, where stretch (10% strain, 0.15Hz) is applied in the presence of endothelium, with the respective log2 fold change and p values. **[F]** Volcano plot of the differentially expressed genes comparing epithelial cells on day 5 of culture, where stretch (10% strain, 0.15Hz) is applied in the presence or absence of endothelium. Genes upregulated and downregulated in the presence of endothelium are shown as green and red respectively. **[G]** Heatmap representation of the 1000 most variant genes. Samples cultured in the presence of endothelium are clustered separately from all other conditions. n=3-4 chips/condition. **[H]** Principal Component Analysis (PCA) shows that the presence of endothelium clearly separates the epithelial transcriptome of the Chips cultured with and without endothelium. n=3-4 Chips/condition.

**Supplementary Figure 2.**
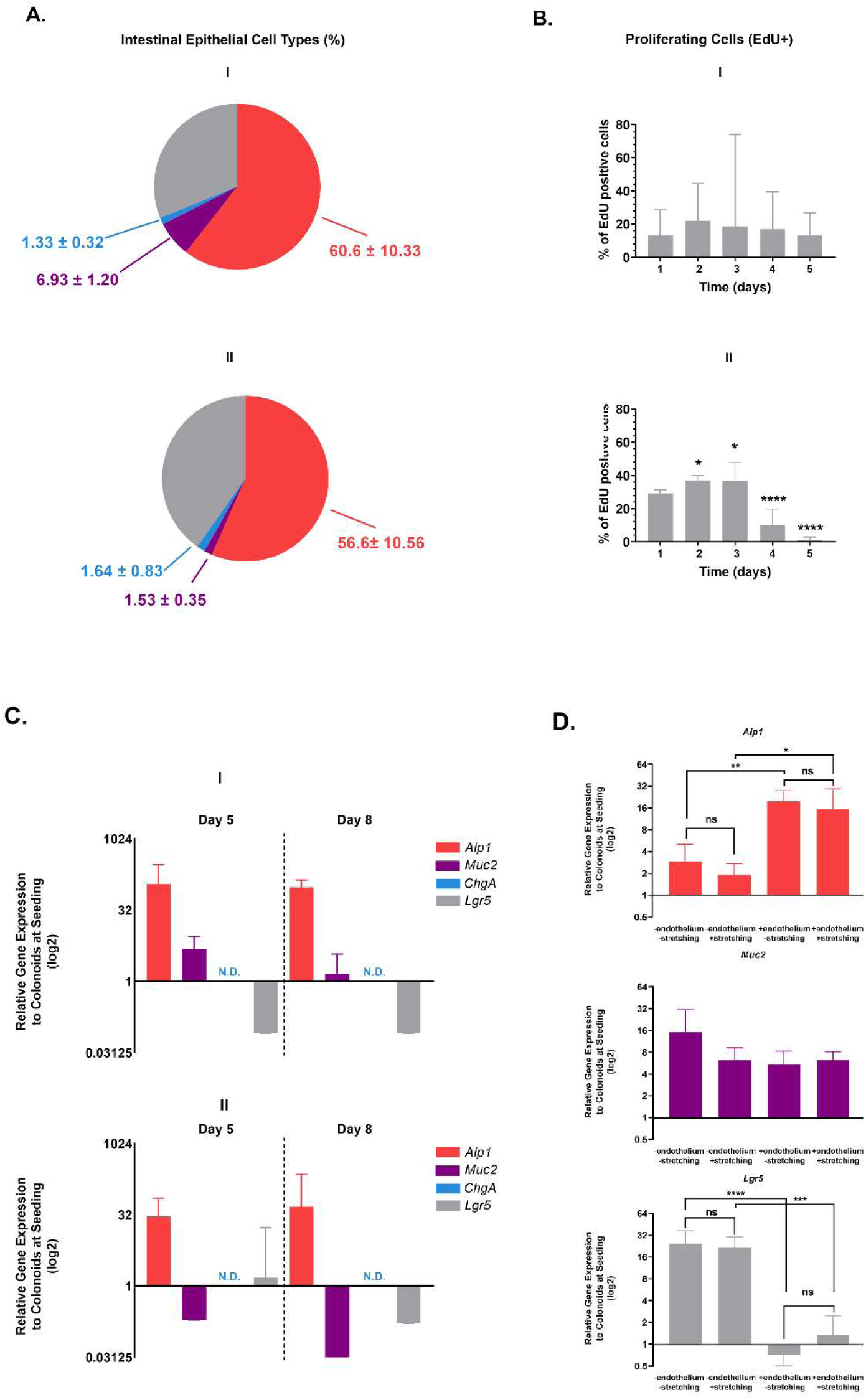
**[A]** Pie charts plot the percentage of the 3 major colonic epithelial cell types. The total number of cells is evaluated based on the DAPI staining. The quantification was performed across 5 different fields of view in 3 individual Chips for each cell type and donor. Values are presented as mean±SD. **[B]** Quantification of the proliferating cells abundance based on EdU staining. The total number of cells is evaluated based on the DAPI staining. The quantification was performed across 3 different fields of view in 3 individual chips for each day and donor. Percentages are plotted as mean±95% CI. The abundance of the proliferating cells is significantly decreased on Day 4 and 5 of the culture. n=3 chips/condition, One-way ANOVA, Tukey’s post hoc test, *: p<0.05, ****: p<0.0001. **[C]** Identification of each epithelial cell type by qPCR for the Alkaline Phosphatase, (Alk1), Mucin 2 (Muc2), Chromogranin A (ChgA), Leukine-rich-repeat-containing G-protein coupled receptor 5 (Lgr5) and polycomb complex protein BMI-1 (Bmi1) genes respectively. The relative gene expression values (RGE) of each gene at the respective timepoints (Day 5 and Day 8) are normalized to colonoids at seeding. 18S has been acquired as the house keeping gene. Values are plotted as mean±95% CI on a log2 scale. A donor dependent expression is observed for the mature enterocytes and cycling stem cells driving genes. **[D]** Identification of each epithelial cell type by qPCR for the Alkaline Phosphatase, (Alk1), Mucin 2 (Muc2), Chromogranin A (ChgA) and Leukine-rich-repeat-containing G-protein coupled receptor 5 (Lgr5) on Day 5 of culture in the presence of stretch and/ or endothelium. Endothelium significantly enhances the gene expression of the absorptive enterocytes’ marker Alp1 and decrease the expression of the cycling stem cells driving gene Lgr5. n=4 chips/condition, One-way ANOVA, Tukey’s post hoc test, ns: p>0.05, *: p<0.05, **: p<0.01, ***: p<0.001, ****: p<0.00001.

**Supplementary Figure 3.**
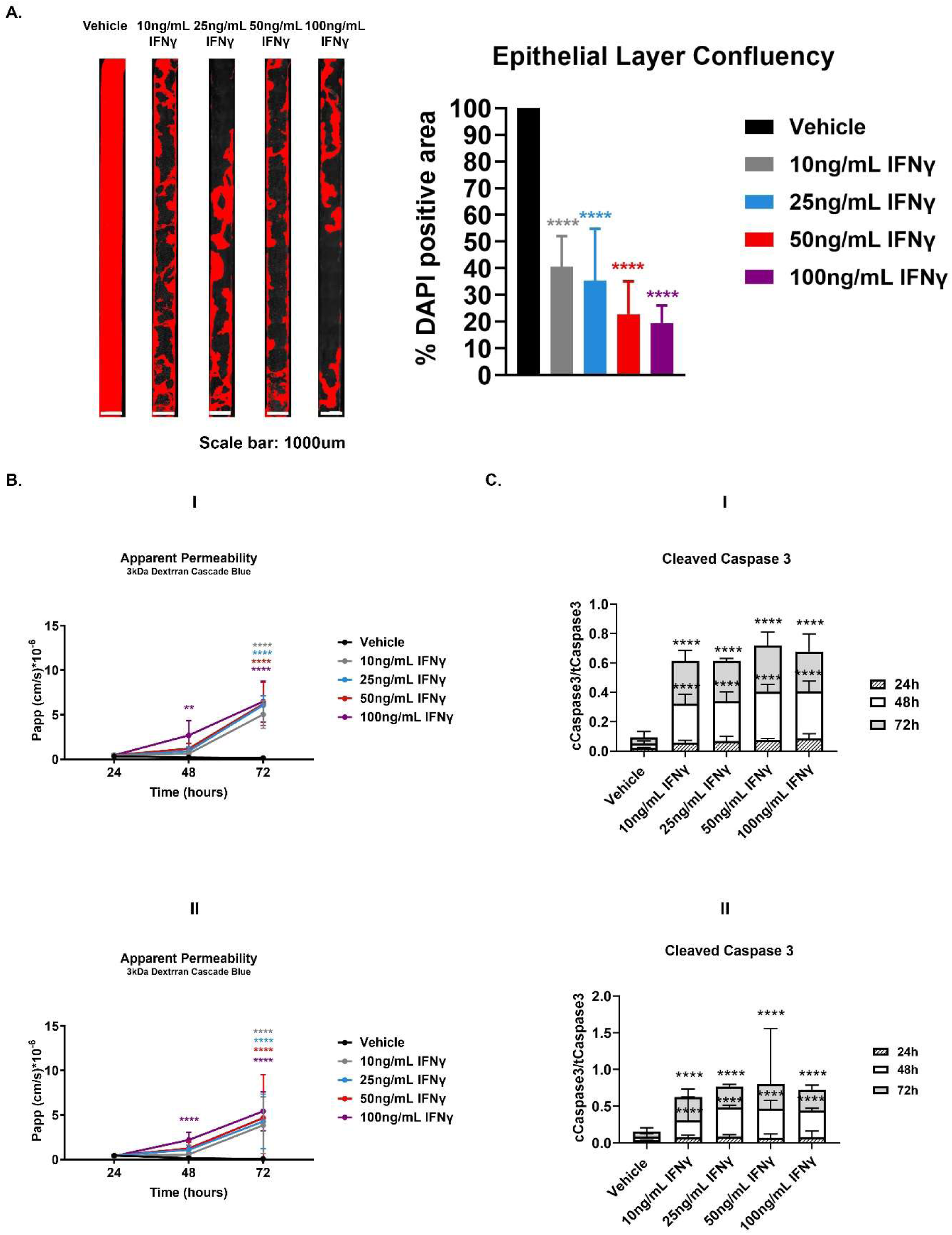
**[A]** Representative tile fluorescent images highlighting the area of the Colon Intestine-Chip covered with epithelial cells, determined by positive DAPI staining (red), 48 h post-stimulation with different concentrations of IFNγ. Scale bar: 1000 μm. The epithelial layer confluency is expressed as a percentage of the DAPI positive area over the total area of the Chip. A significant and concentration dependent decrease of the epithelial layer confluency is observed 48 h post-treatment. n=3 Chips/condition, mean±95% CI, One-way ANOVA, Tukey’s post hoc test, ****: p<0.0001. **[B]** Apparent Permeability (P_app_) of the colonic epithelial cells to 3kDa Dextran Cascade blue, over the course of 72 h basolateral stimulation with IFNγ. A dose and time dependent disruption of the barrier integrity is observed for each donor. Data shown correspond to one representative out of two independent experiments. n= 3-9 Chips/condition, mean±95% CI, Two-way ANOVA, Tukey’s post hoc test, **: p<0.01, ****: p<0.0001. Scale bar: 50μm. **[C]** Quantification of the cleaved over the total Caspase 3 in cell lysates from the epithelial channel of the Colon Intestine-Chip. A significant and concentration dependent increase in the levels of cleaved Caspase 3 is observed 48 h and 72 h post-stimulation for both donors. n= 2-3 Chips/ condition mean±95% CI, Two-way ANOVA, Tukey’s post hoc test, ****: p<0.0001.

**Supplementary Table 1.**
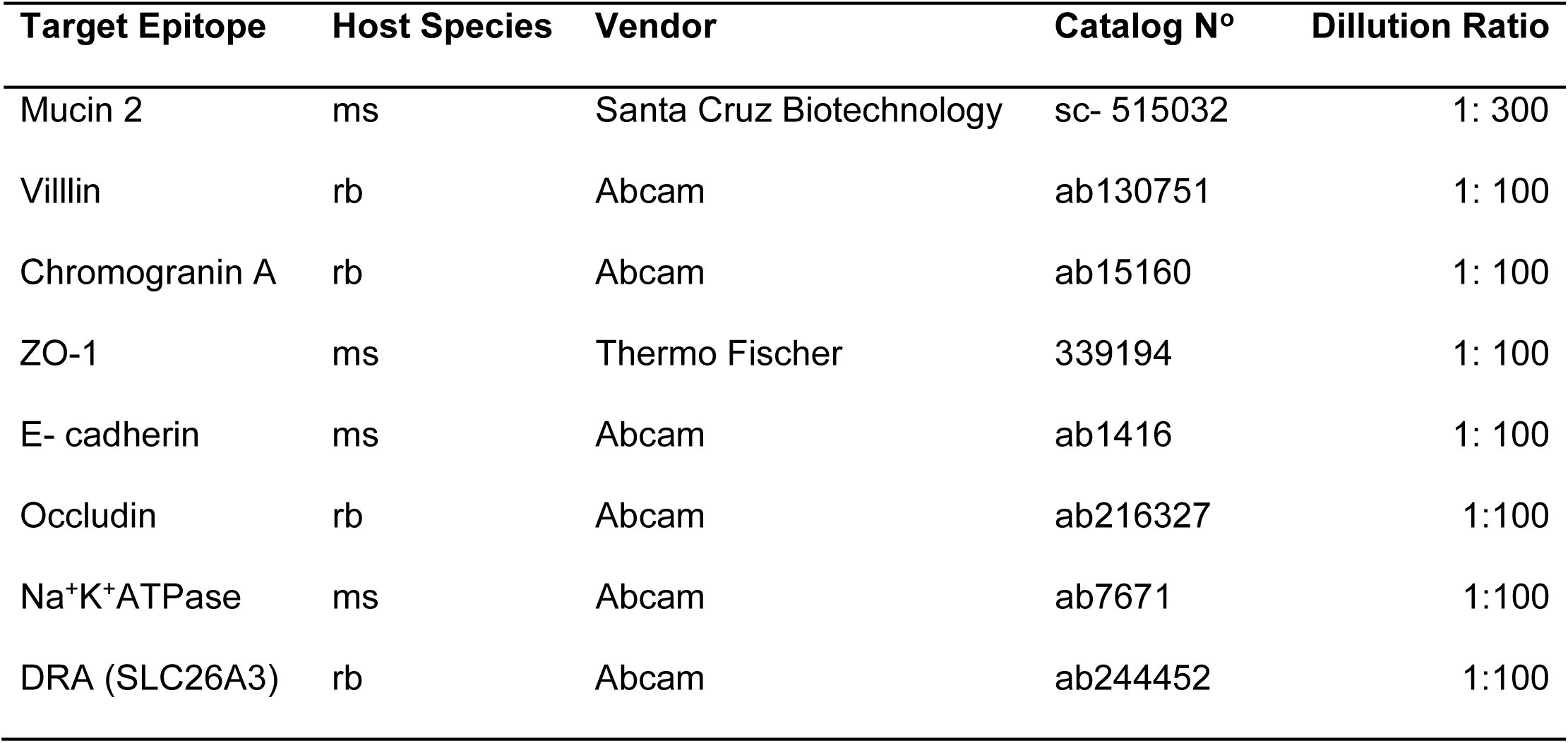
List of antibodies used in immunofluorescence

**Supplementary Table 2.**
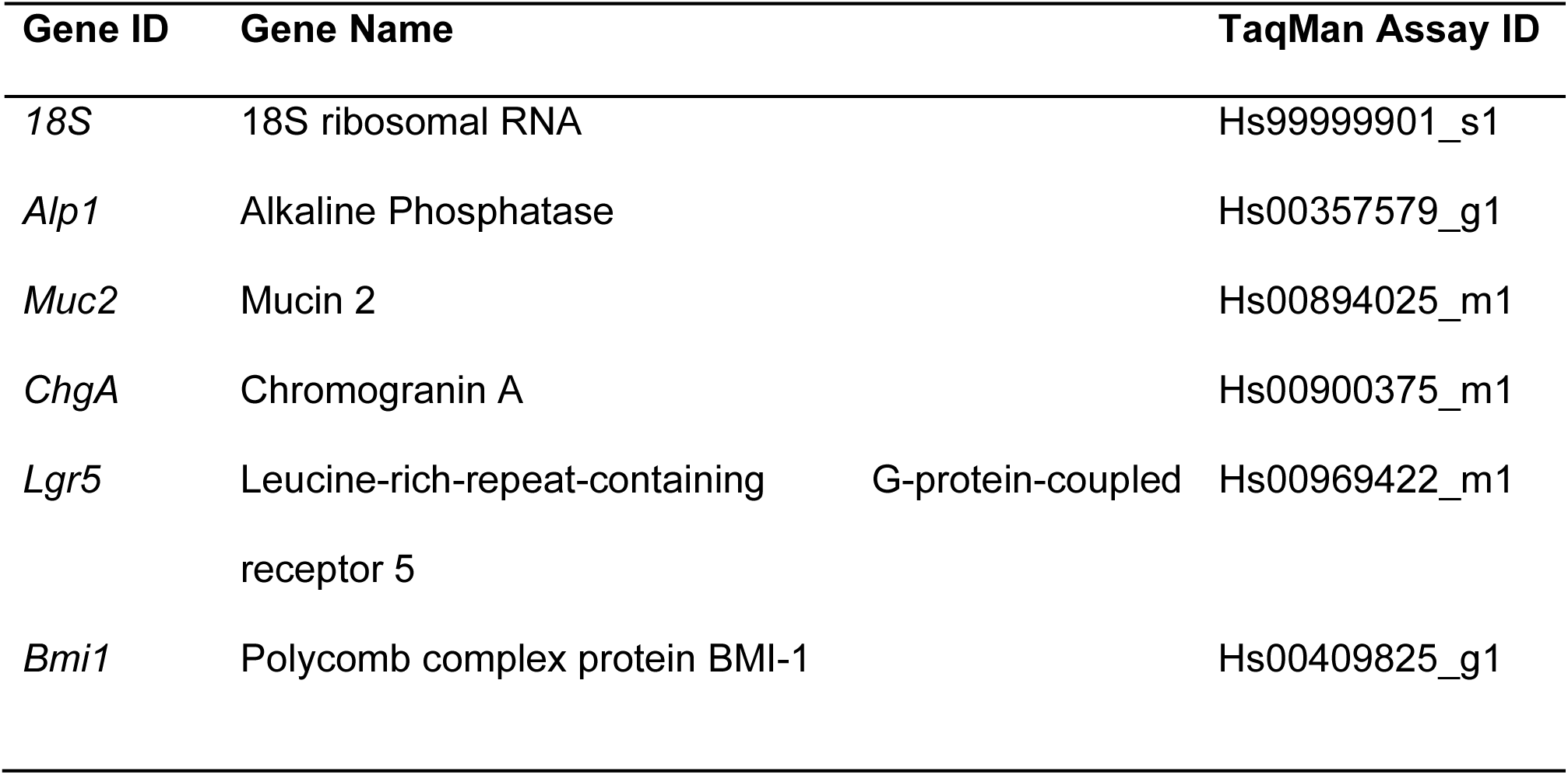
List of human TaqMan Gene Expression Assays used in qRT-PCR

**Supplementary Table 3.** List of samples where RNA sequencing was performed.

| **Timepoint** | **Experimental Group** | **N° of Replicates** |
| --- | --- | --- |
| Day 0 | Colonoids | 3 |
| Day 5 | -endothelium/ -stretch | 4 |
|  | -endothelium/ +stretch | 4 |
|  | +endothelium/ -stretch | 4 |
|  | +endothelium/ +stretch | 6 |
| Day 8 | -endothelium/ -stretch | 4 |
|  | -endothelium/ +stretch | 4 |
|  | +endothelium/ -stretch | 4 |
|  | +endothelium/ +stretch | 3 |

